# A metagenomic view of novel microbial and metabolic diversity found within the deep terrestrial biosphere

**DOI:** 10.1101/2021.05.06.442964

**Authors:** Lily Momper, Caitlin P. Casar, Magdalena R. Osburn

## Abstract

The deep terrestrial subsurface is a large and diverse microbial habitat and a vast repository of biomass. However, in relation to its size and physical heterogeneity we have limited understanding of taxonomic and metabolic diversity in this realm. Here we present a detailed metagenomic analysis of samples from the Deep Mine Microbial Observatory (DeMMO) spanning depths from the surface to 1.5 km deep in the crust. From these eight geochemically and spatially distinct fluid samples we reconstructed ∼600 metagenome assembled genomes (MAGs), representing 50 distinct phyla and including 18 candidate phyla. These novel clades include many members of the Patescibacteria superphylum and two new MAGs from candidate phylum OLB16, a phylum originally identified in DeMMO fluids and for which only one other MAG is currently available. We find that microbes spanning this expansive phylogenetic diversity and physical space are often capable of numerous dissimilatory energy metabolisms and are poised to take advantage of nutrients as they become available in relatively isolated fracture fluids. This metagenomic dataset is contextualized within a four-year geochemical and 16S rRNA time series, adding another invaluable piece to our knowledge of deep subsurface microbial ecology.

## INTRODUCTION

Earth’s deep biosphere is a vast repository of microbial biomass. Although the size of this biosphere has been continually refined, the most recent estimates now suggest that it is one of the largest biomes on the planet, ranging from 15-23 Pg to perhaps as much as 60 Pg of carbon sequestered in resident microbes (Whitman *et al*., 1998; McMahon and Parnell, 2013; Magnabosco *et al*., 2018; Bar-On *et al*., 2018; Flemming and Wuertz, 2019). The deep subsurface biosphere (DSB) found underlying the continents or terrestrial biosphere, has emerged as a dynamic, populated, metabolically-active environment, affecting carbon storage and global elemental cycles (Barry et al., 2019; Magnabosco et al., 2018; Flemming and Wuertz, 2019). Although some DSB environments exhibit exceptionally low diversity (Chivian et al., 2008), many of those studied harbor an abundant and diverse microbiome (e.g., Baker et al, 2016; Chivian et al, 2008; Dong et al, 2014; Lau et al, 2014; Magnabosco et al, 2015; Magnabosco et al., 2018; Nyyssönen et al, 2014; Rinke et al, 2013). Furthermore, newly identified subsurface microbial lineages are continually and commonly implicated in major geochemical cycles, highlighting the importance of uncultivated groups in understanding those cycles (Baker et al., 2016; Rasigraf et al., 2014). However, the vast, difficult to access, and heterogeneous nature of the terrestrial DSB complicate efforts to study microbially mediated processes therein. Further confounding our understanding of these processes is the growing appreciation that a high proportion of subsurface microbes have yet to be cultured, known only by culture-independent genetic evidence and colloquially referred to as ‘microbial dark matter’ (Castelle et al., 2015; Momper et al., 2017; Rinke et al., 2013; Wrighton et al., 2012).

Microbial observatories are portals into the deep subsurface established to monitor subsurface geochemistry and microbial ecology across the world, including in Sweden, Finland, Canada, Switzerland, and the United States (Cardace et al., 2013; Osburn et al., 2019; Pedersen, 1996; Pedersen et al., 2014; Stroes-Gascoyne et al., 2007). The advent of high-throughput shotgun DNA sequencing methods was a pivotal technological advance towards understanding the microbial diversity of the deep terrestrial subsurface, reducing primer bias and capturing the full breadth of *in situ* diversity. Recent applications of these techniques have revealed that shallow and deep subsurface environments harbor a vast diversity of subsurface taxa particularly in uncultivated, candidate, groups and that spatial separation of taxa occurs with depth and between aquifers (Anantharaman *et al*., 2016; Jungbluth *et al*., 2016; Probst *et al*., 2017; Momper et al., 2017a; Momper et al., 2017b; Probst *et al*., 2018). Key genomic adaptations discovered within deep intraterrestrials include widespread capacity to fix carbon, particularly with the Wood Lungdal pathway (Lau et al., 2016; Magnabosco et al., 2016; Momper et al., 2017) and a dichotomy of small, ultra-streamlined genomes and larger, bulky genomes with diverse metabolic capabilities (Anantharaman et al., 2016; Jungbluth et al., 2016; Jungbluth et al., 2017; Lau *et al*., 2016). Current comprehensive reviews of microbial life on and within Earth integrate information from these vast sequencing datasets, estimate the diversity and abundance of the DSB as a whole, and further underscore subsurface microbes’ impact on global biogeochemical processes (Flemming and Wuertz, 2019; Magnabosco et al., 2018). However, to date these datasets are heavily focused on shallow subsurface realms, or individual locations within a single very deep site, often without geochemical context, leaving a gap in the range of depths and environments for which we have a clear genomic and metabolic understanding.

In this study we use whole metagenome sequencing methods to investigate microbes from the Deep Mine Microbial Observatory- DeMMO-(Osburn et al., 2019) located in the Sanford Underground Research Facility (SURF) in Lead, South Dakota, USA. We draw on a four-year geochemical and 16S rRNA tag sequencing dataset collected at DeMMO (Osburn et al., 2019, 2020) to integrate microbial, geochemical, and geological observations to expand our genomic interpretation to subsurface microbial processes. Notably, within this metagenomic dataset, we have near complete metagenome assembled genomes (MAGs) from a rich array of previously uncultured bacteria. We analyze these MAGs to query and characterize metabolic capabilities of microbial communities at DeMMO and use phylogeny of binned MAGs to tie those metabolic capabilities to responsible taxa. We focus our analysis on MAGs from uncultivated groups, especially those for which ours are the most complete genomes to date, as this is the primary mechanism to understand the metabolic role of microbial dark matter (MDM) in the terrestrial DSB.

## MATERIALS AND METHODS

### Field sampling

All samples for sequencing and corresponding geochemical data were collected from DeMMO borehole fluids and controls (mine service water used for lubricant during borehole drilling and an overlying freshwater stream – Whitewood Creek) between April 16-19, 2018. Microbial cells were filtered from 10 L of fluid onto 47 mm, 0.1 µm Supor filters (Pall Corporation, Port Washington, NY, USA), frozen immediately on dry ice and stored frozen at - 80°C. Detailed descriptions of the host geology, borehole characteristics, the establishment of the DeMMO, and methods for geochemical analyses can be found in Osburn et al. (2014) and Osburn et al. (2019). Physical and geochemical characteristics of fluids at the time of sampling are provided in Table 1.

**Table 1.** Measurements of geochemical constituents collected concomitantly with biologcial samples for DNA extraction and sequencing ^*^Dissolved gas concentrations were not measured on the day of collection in April of 2018. We have used an averaged value from measurements taken December 2015-December 2019 in corresponding boreholes.

| Site | Date | Flow rate<br>mL/min | Temp<br>(°C) | Conductivity<br>(μS) | TDS<br>ppm | pH | ORP<br>(mV) | NO<sub>3</sub><sup>-</sup><br>mg/L | NH<sub>4</sub><sup>+</sup><br>mg/L |
| --- | --- | --- | --- | --- | --- | --- | --- | --- | --- |
| DeMMO 1 | 4/18/2018 | 3000 | 10.5 | 1045 | 756 | 5.83 | -68 | 0.3 | 0.06 |
| DeMMO 2 | 4/18/2018 | 300 | 12 | 609.4 | 430.4 | 7.44 | -100 | 0.3 | 0.03 |
| DeMMO 3 | 4/17/2018 | 4600 | 16.1 | 3082 | 2349 | 7.13 | -61 | 0.3 | 0.21 |
| DeMMO 4 | 4/19/2018 | 300 | 22.2 | 1781 | 1302 | 7.46 | -200 | 1.1 | 1.36 |
| DeMMO 5 | 4/16/2018 | 15600 | 31.6 | 1534 | 1103 | 8.45 | -149 | 0.7 | 0.48 |
| DeMMO 6 | 4/16/2018 | 0 | 20.1 | 7933 | 6602 | 6.61 | -198 | 0.3 | 0 |

\*Dissolved gas concentrations were not measured on the day of collection in April of 2018. We have used an av

xtraction and sequencing
| Fe<sup>2+</sup> | Total sulfide | DO | SO<sub>4</sub><sup>2-</sup> | Total Fe | Total Mn | DOC | DIC | H<sub>2</sub> * | CH<sub>4</sub> * |
| --- | --- | --- | --- | --- | --- | --- | --- | --- | --- |
| mg/L | μg/L | mg/L | mg/L | mg/L | mg/L | mg/L | mM | mg/L | mg/L |
| 2.32 | 6 | 0.025 | 393 | 6.08 | 0.288 | 0.468 | 3.81 | 1.19E-07 | 7.75E-06 |
| 0.31 | 0 | 0.054 | 84.8 | 0.35 | 0.052 | 0.393 | 4.02 | 1.42E-07 | 5.85E-06 |
| 2.86 | 5 | 0.029 | 1800 | 3.3 | 0.297 | 0.25 | 9.46 | 2.68E-07 | 6.88E-05 |
| 0 | 582 | 0.031 | 315 | 0.03 | 0.005 | 0.244 | 11.31 | 2.16E-07 | 0.000719 |
| 0 | 254 | 0.015 | 186 | 0 | 0 | 0.316 | 10.96 | 1.98E-07 | 0.000346 |
| 1.47 | 13 | 0 | 4110 | 2.44 | 0.375 | 0.286 | 1.86 | 3.05E-07 | 0.005041 |

eraged value from measurements taken December 2015-December 2019 in corresponding boreholes.
| CO<sub>2</sub> * | CO * |
| --- | --- |
| mg/L | mg/L |
| 0.022831 | 5.56E-06 |
| 0.017753 | 4.94E-06 |
| 0.091982 | 8.19E-06 |
| 0.013094 | 3.54E-06 |
| 0.00477 | 4.14E-06 |
| 0.002487 | 2.52E-06 |

### DNA extraction and sequencing

Whole genomic DNA was extracted using a modified phenol-chloroform method with ethanol precipitation as described in Osburn et al. (2014). Library preparation, pooling, quality control and sequencing were performed at the Environmental Sample Preparation and Sequencing Facility at Argonne National Laboratory, Lemont, IL, USA. Sequencing was performed on an Illumina HiSeq2500 platform, resulting in ∼150 bp paired end reads.

### De novo assembly, read mapping and generation of metagenome assembled genomes

Trimming of paired end reads was performed using Trimmomatic 0.36 with default parameters and a minimum sequence length of 36 base pairs (Bolger et al, 2014). Reads were assembled using the MEGAHIT assembly algorithm (Li et al., 2015) with a 1,000 bp minimum contig length. Coverage depth information was then generated for scaffolds greater than 1,000 base pairs by mapping the 150 base pair paired-end reads of each sample to its respective assembly using the Burrows-Wheeler Alignment tool, BWA version 0.7.15 (Li and Durbin, 2009) with the BWA default parameters. SAMtools v0.1.17 (Li et al, 2009) was then used to convert files to binary format for downstream analysis.

Metagenome assembled genomes (MAGs) were generated using MetaBAT2, which relies on sequence composition, differential coverage and read-pair linkage (Kang et al., 2019). MAG completeness (reported as percentage of the set of single copy marker genes present) and contamination (calculated as multiple occurrences of a single copy marker gene) were calculated using the “lineage_wf” workflow in the CheckM pipeline (Parks et al, 2015). MAGs that contained >10% contaminating single copy genes were manually refined and curated using the Anvi’o program (Eren et al., 2015). MAG completeness and contamination were subsequently re-calculated using five different standard marker gene suites (Alneberg et al, 2014; Campbell et al, 2013; Creevey et al, 2011; Dupont et al, 2012; Wu & Scott, 2012). MAGs with >10% contamination after curation were removed. Here, we have included all MAGs for which taxonomy could be assigned, even if those with low completeness (14-30%). Lowering the completeness threshold allowed greater retention of DNA generated from each sample and wider taxonomic representation. Because our analysis relies on gene presence, not absence, the inclusion of these less complete MAGs does not compromise our analyses.

### Assignment of putative taxonomies and ribosomal protein tree construction

MAGs were assigned taxonomic identities first according to their placement in a phylogenomic tree using the “tree” command in CheckM (Parks et al, 2015) and refined using the Genome Taxonomy Database and the associated GTDB-Tk toolkit (Chaumeil, et al. 2019; Parks et al., 2018; Parks et al., 2020 https://github.com/Ecogenomics/GtdbTk). A ribosomal protein tree was generated using GToTree (Lee, 2019) and RAxML (Stamatakis, 2014). MAGs were queried for the 15 syntenic and phylogenetically informative ribosomal proteins identified previously (Hug et al., 2016) using HMMER v3.1b1 (Eddy, 2011). If multiple single copy target genes were identified in a single MAG, that particular gene was excluded from the alignment. Ribosomal protein sequences were aligned independently using MUSCLE v. 3.8.31 (Edgar, 2004). Automated trimming of alignments was performed with Trimal (Gutıerrez et al., 2009) and individual alignments from a given MAG were concatenated. MAGs were removed from the final alignment if <40% of the 15 queried ribosomal proteins were identified. 1,673 reference genomes were downloaded from the NCBI database and processed using the same methods to produce a concatenated ribosomal protein alignment for each genome. A maximum likelihood tree generated from this alignment was constructed using RAxML v. 8.2.12 (Stamatakis, 2014) under the LG plus gamma model of evolution (PROTGAMMALG in the RAxML model section), with 1000 bootstrap replicates.

### Metabolic pathway analysis

Gene calling and metabolic pathway identification in MAGs was performed using METABOLIC (METabolic And BiogeOchemistry anaLyses In miCrobes) Version 3.0 (Zhou et al., 2019). METABOLIC uses a combination of KOFAM, Pfam, TIGRfam, MEROPs, dbCan2, and customized Hidden Markov Models (HMMs) to identify and classify coding sequences. If >75% of the genes for a given pathway were identified in a single MAG, that pathway was considered present and ‘complete’, regardless of that MAG’s estimated genome completeness.

## RESULTS

### Metagenome assembled genomes and phylogenetic identification

A total of 581 MAGs were recovered from the eight assembled metagenomes. Genome statistics including MAG ID, NCBI taxon ID, completeness and contamination are listed in Supplementary Data File 1. Of all these 581, 81 were >90%, 99 were 70-89% and 154 were 50-69% complete (Fig. 1). Of the less complete genomes (< 50% complete), more than a quarter (56 of 211) belong to the Candidate Phyla Radiation (CPR). This group is known to contain streamlined genomes often lacking the full suite of ‘essential’ single copy marker genes, and therefore is not necessarily a reflection of poor assembly (Nelson and Stegen, 2015; Castelle and Banfield, 2018). Similarly, Whitewood Creek (WC), the stream water control, contained the greatest proportion of incomplete genomes (∼75% were <50% complete) and the highest proportion of CPR MAGs at 47% (28 MAGs) (Fig. 2a and c).

**Figure 1.**
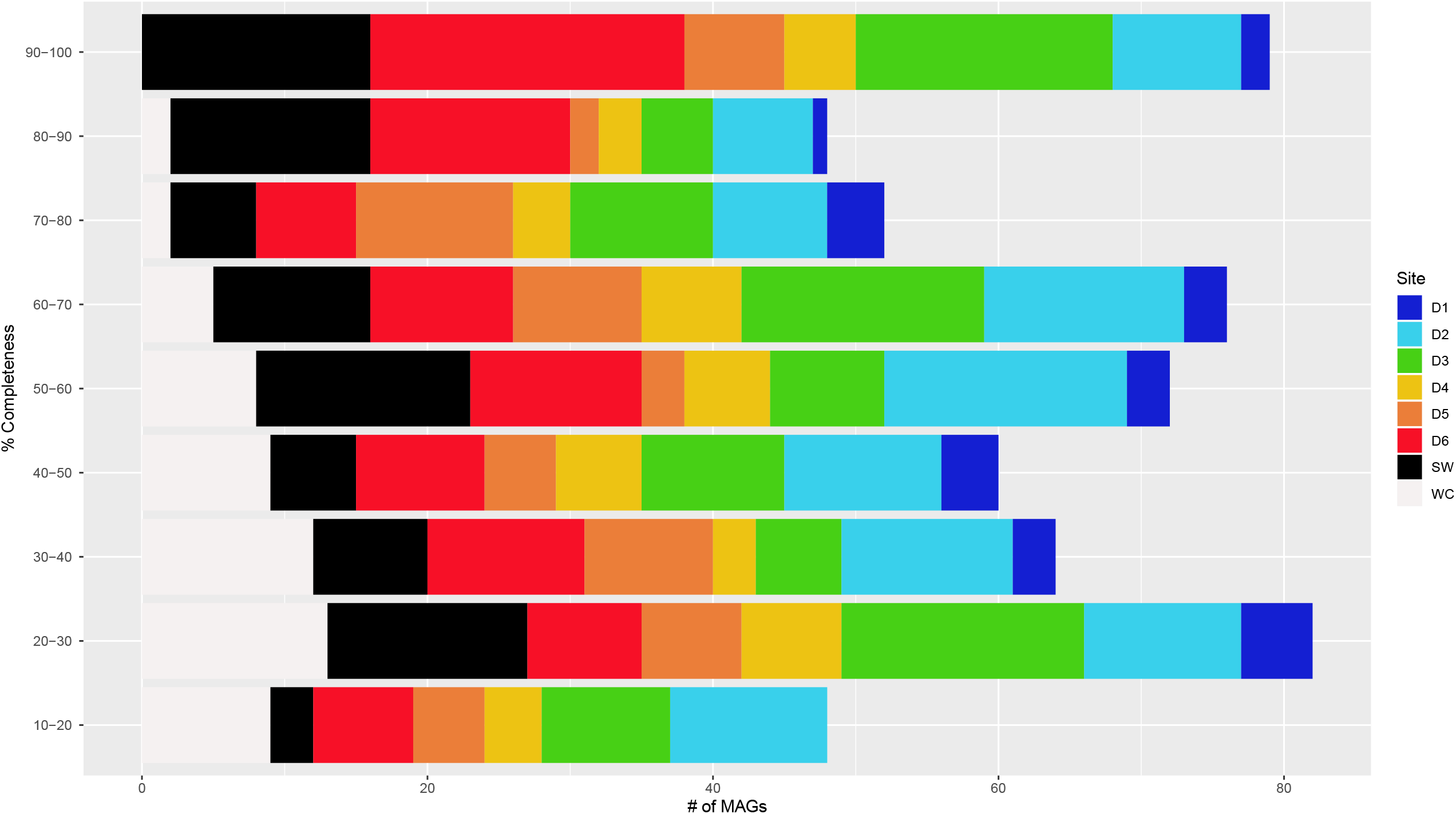
Statistical breakdown of MAG completeness for all 582 genomes across eight samples.

**Figure 2.**
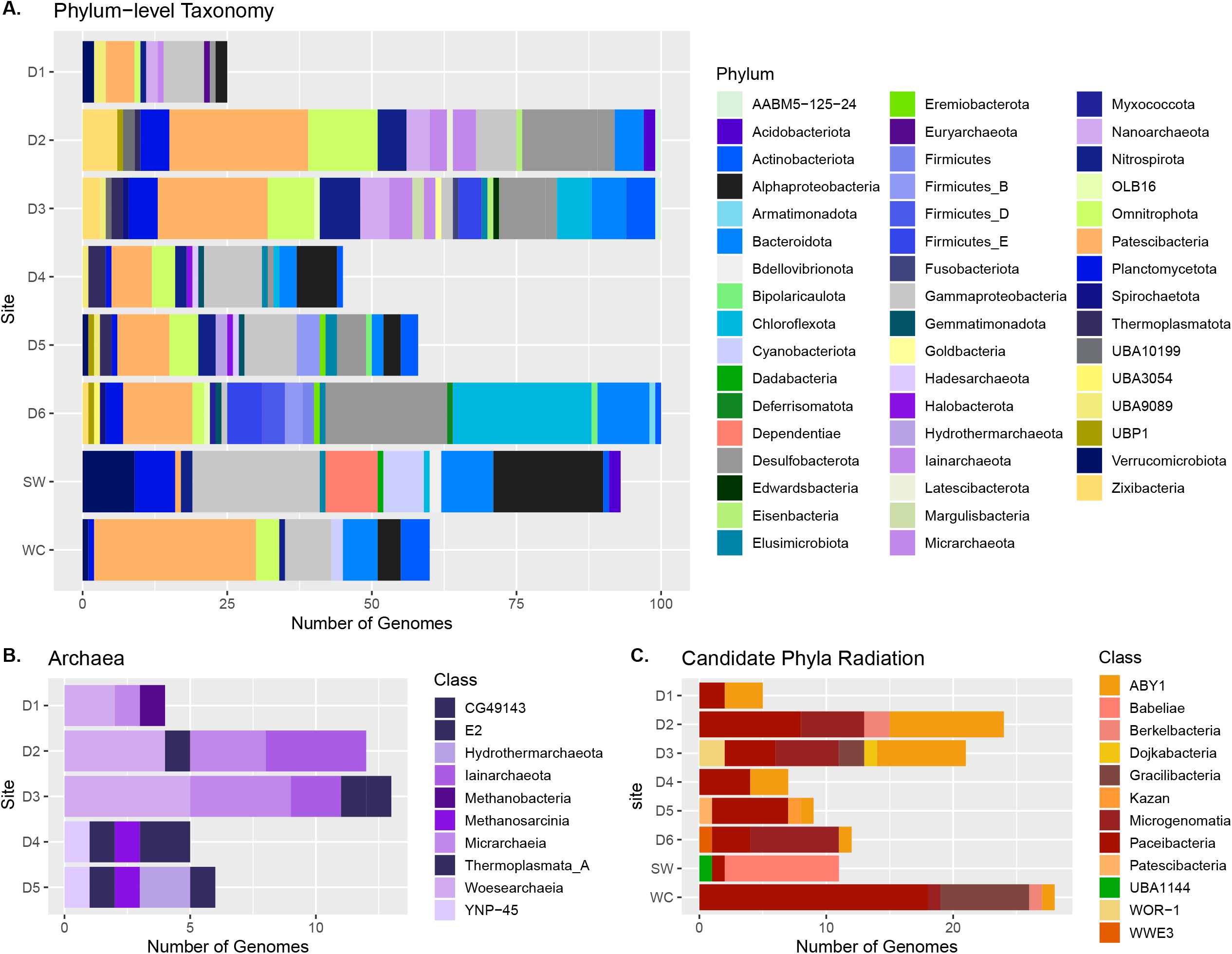
Taxonomic grouping of all MAGs reconstructed in this study, at the phylum and class levels. A) Phylum level taxonomic assignments for MAGs in terms of relative abundance per sample. B) Classes of Archaea identified in fluids. C) MAGs from the Candidate Phyla Radiation (CPR) identified in this study.

Of the six DeMMO samples, hereafter called D1-D6, the fewest MAGs were recovered from D1 (25) and the most from D6 (100) (Fig. 1 and Fig. 2). The controls, hereafter referred to as Service Water (SW) and WC, differ from the DeMMO samples in that 1) they lack Archaea 2) the WC fluid was enriched in the CPR (Fig. 2c) and 3) the SW contained more than twice the number of Proteobacterial MAGs of any other sample (Fig. 2d). Phylum level taxonomic assignments for MAGs can be found in Figure 2a, class break down of the Patescibacteria and Archaea in Figure 2b and 2c, and assignment to the lowest confident taxonomic level in Supplementary Data File 1. MAGs are spread across the phylogenetic tree (Fig. 3). Notably, this dataset is enriched in members of the CPR and uncultivated groups (candidate phyla and classes) compared to groups with cultivated members (Fig. 2 and 3). Currently there are 14 recognized classes within the Patescibacteria (aka CPR) (Parks et al., 2018), 12 of which are found here (Fig. 2c). Uncultured and candidate phyla are present in all sites, including controls and DeMMO boreholes, totaling more than 20 such groups (Fig. 2).

**Figure 3.**
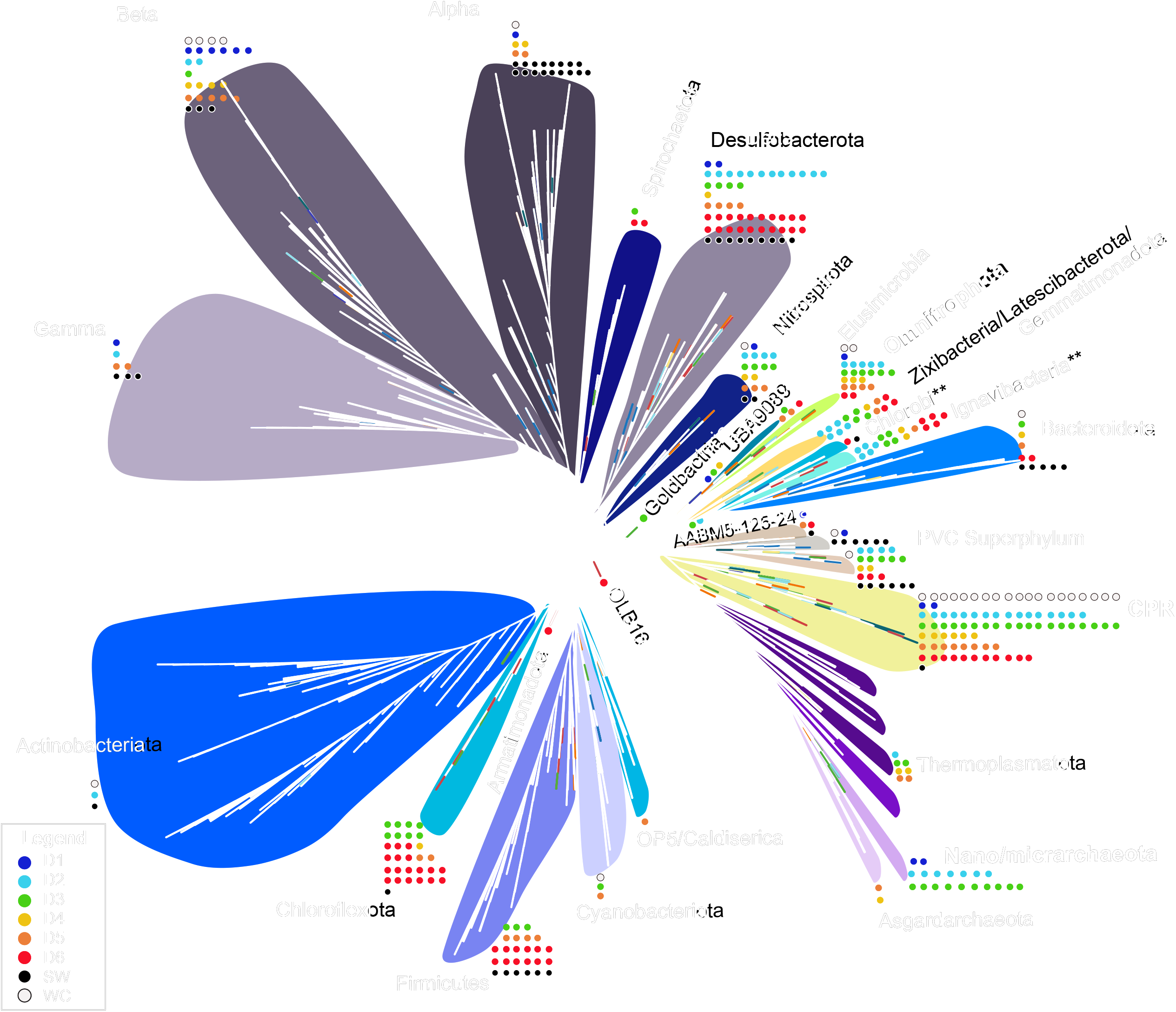
Concatenated ribosomal protein tree containing all MAGs for which at least 40% of target ribosomal proteins could be identified. Phyla with ** (Chlorobi and Ignavibacteria) have traditionally been considered separate phyla within the FCB superphylum. The GTDBTk toolkit has included them in the Bacteroidota phylum. We have kept them separate in the ribosomal protein tree for clarity and because it is not yet widely accepted that they should be classified in the same phylum.

### Metabolic capabilities in DeMMO fluids planktonic communities

Our objective in this analysis is to present an overview of the metabolic potential of DeMMO genomes over the represented spatial and geochemical landscape and to attribute this potential to specific taxonomic groups. To this end, we present genes for prominent energy metabolisms at all sites including WC and SW to contrast surface waters to subsurface fluids. Genes which are diagnostic for a variety of dissimilatory metabolisms were queried in MAGs and those of most interest are displayed in Fig. 4 (complete gene name, Pfam/TIGRfam are in Supplementary Data File 2). A complete list of all gene hits for all MAGs is presented in Supplementary Data File 3.

**Figure 4.**
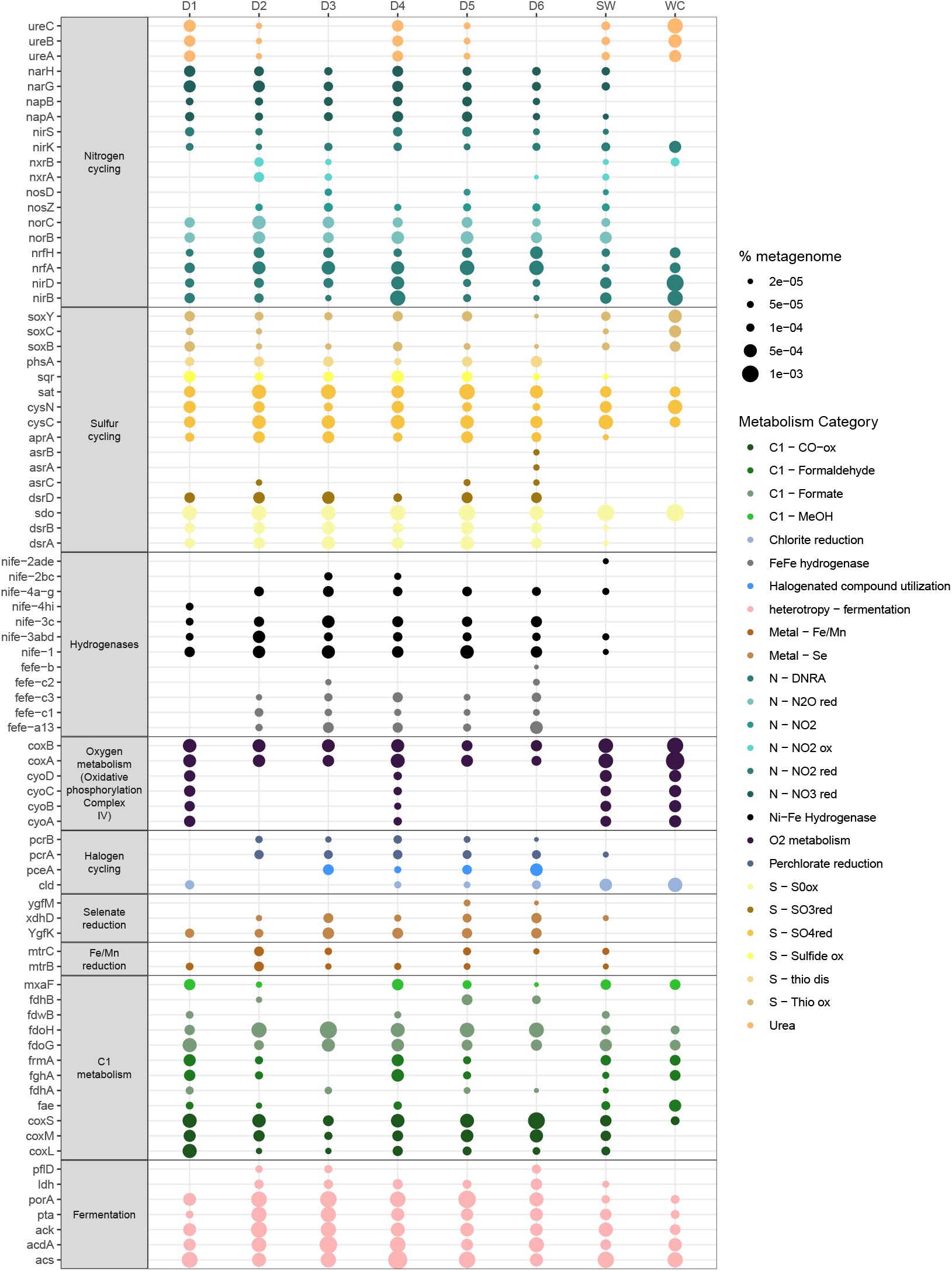
Functional gene annotations indicative of potential energy-yielding metabolisms in MAGs across all eight fluid samples. Canonical genes for common electron donors and acceptors were queried in all MAGs binned from DeMMO fluids and two control samples.

### Nitrogen

Nitrogen is present in many oxidation states (−3 to +5) which are transformed through a variety of microbially mediated redox reactions. In respiratory denitrification, nitrate (NO_3_^-^), nitrite (NO_2_^-^), nitric oxide (NO) and nitrous oxide (N_2_O) are sequentially reduced to dinitrogen gas (N_2_), with each step catalyzed by one or more metalloenzymes: *nar, nir, nor* and *nos* catalyzing each step (Zumft 1997). The potential for nitrate reduction is present all samples, with the intracellularly (*nar*) mediated reduction being relatively more abundant than the periplasmic (*nap*) form (Fig. 4). Microbial nitrite reduction (NO_2_^-^ to NO) is catalyzed by two related respiratory periplasmic enzymes: the *nirK-*encoded copper nitrite reductase or the *nirS-*encoded cytochrome *cd*_*1*_ (Berks et al., 1995; Brittain et al., 1992; Green et al., 2010). These genes are relatively scarce compared to those that catalyze nitrate reduction, however they were identified in all samples with the exception of *nirS* in D3 MAGs (Fig. 4). The metabolic potential for the final step in denitrification varies across samples as *nosD* was identified in MAGs from D3 and 5, but not D1, D4 of D6 and the canonical marker gene, *nosZ*, was found in D2-D6, but not D1 (Fig. 4). Dissimilatory nitrate reduction to ammonium (DNRA) bacteria use NrfA as their key enzyme. Assisted by its redox partner NrfH, NrfA catalyzes the six-electron reduction of nitrite to ammonium (Simon, 2002; Einsle, 2011; Simon and Klotz, 2012). Genes for both subunits are highly abundant in all DeMMO samples (Fig. 4). The *nxrA* and *nxrB* genes are powerful functional and phylogenetic markers to detect and identify uncultured nitrite oxidizing bacteria (Anantharaman et al., 2016; Crane et al., 1995; Daniels et al., 1986) and these genes are most prominent in D2, with lesser detection in D3, D5 and D6 (Fig. 4).

### Sulfur

Sulfur is a versatile element and can be metabolically transformed between eight oxidation states (−2 to +6). Canonical genes involved in sulfite (SO_2_^-^), sulfate (SO_4_^3-^) and thiosulfate (S_2_O_3_^2-^) reduction (*sat, asr*, and *phs*, respectively) were queried in all MAGs. *Sat* was highly abundant across all DeMMO samples, while *phsA* was less common, and the complete set of *asr* subunits (*A, B* and *C*) were identified exclusively in D6 (Fig. 4). Sulfur species can also be used as electron donors. Genes involved in sulfur oxidation, *sqr* and *sdo*, were identified in all DeMMO samples and *sdo* was relatively abundant (Fig. 4). SQR is known to be used for sulfide (H_2_S) detoxification and therefore is not necessarily indicative of dissimilatory processes, but regardless its presence indicates the potential to transform H_2_S to elemental sulfur (S^0^) or sulfate (SO_4_^2-^). Microbially mediated S transformation in the environment can be quite complex as many enzymes that transform sulfur can use it as a terminal electron acceptor (TEA) or electron donor, depending on environmental conditions. Among these are dissimilatory sulfite reductase (*dsr*), adenosine-5′-phosphosulfate (*aprA*) and the Sox gene cluster (*sox*). The *dsr* genes are the most common across all sites, followed by *aprA*. Sox genes were far less common, most abundant in D1, but *soxC* was not identified at all in D3 and D6 (Fig. 4). *DsrD*, the small subunit of dissimilatory sulfite reductase that is essential for sulfate reduction, corresponds to the abundance of *dsrAB* in D1-D3 and D6, but is slightly less abundant in D4 and D5 (Fig. 4).

### Hydrogen and Oxygen

Hydrogen gas is present in DeMMO fluids (Osburn et al., 2014; Osburn et al., 2019) and can be a powerful electron donor in subsurface environments (Nealson et al., 2005; Takai et al., 2004). We queried a suite of hydrogenases involved in hydrogen-transforming microbial reactions (Sondergaard et al., 2016). Among those hydrogenases, NiFe groups 1, 3ABD and 3C were identified in all six DeMMO fluids and were the most abundant hydrogenases. FeFe group B were least abundant, with identification in only D6. FeFe group C2 and NiFe group 4H were also relatively uncommon, identified only in D3 and D1, respectively. D1 had by far the fewest number and types of hydrogenases whereas D3 and D6 had the greatest.

Aerobic bacteria and archaea use oxygen as their respiratory TEA. Oxygen-respiring genes are most abundant in the control WC and SW samples, followed closely by D1, and D4 (Figure 4). Cytochrome oxidase (Cox) catalyzes the reduction of oxygen to water and is therefore essential for aerobic metabolism (Capaldi, 1990; Chan and Li, 1990; Saraste, 1990; Babcock and Wikstrom, 1992). Subunits *coxA* and *coxB* were found in all eight samples (Figure 4). Another terminal oxidase, the *bo*_3_ oxidase (Cyo), was found in four of the eight samples: D1, D4, WC and SW (Figure 4). In summary, oxygen respiring genes were found in all samples, but the distribution of specific genes differed between sites.

### Metals and halogens

Known genes for iron, manganese and selenite reduction, as well as those involved in halogen cycling, were identified in all samples to varying degrees. Genes indicative of metal reduction, *mtrB, mtrC*, were detected in all DeMMO fluids, relatively most common in D2, but not detected in WC. Putative selenate reduction genes were found in all samples save WC and was highest in D5 and D6 (Fig. 4). The canonical gene for chlorite reduction (*cld*) was particularly abundant in the controls, whereas putative perchlorate reduction (*pcrA* and *pcrB*) was identified in samples D2-D6, but not controls. The gene implicated in reductive halogenation (*pceA*), typically associated with members of the phylum Chloroflexi (specifically the *Dehalococcoidia*), was found in D3-D6, with highest abundances in D3 and D6 (Fig. 4) corresponding to the relative abundance of Chloroflexi.

### Carbon respiration and fixation

C1 carbon metabolisms were investigated owing to their potential importance to subsurface metabolism including carbon monoxide (CO), methanol (CH_3_OH), formate (CHO_2_^-^), and formaldehyde (CH_2_O) oxidation. Genes known to be involved in CO oxidation (*coxLMS*) were abundant in all sites except WC and those used in formate oxidation (*fdhAb, fdoHG, fdwB*) were abundant across all samples (Fig. 4). Genes responsible for formaldehyde oxidation (*fae, fghA, frmA*) were variable across samples: they were not present in D3 or D6, at low abundance in D2 and D5, and most abundant in D1, D4 and controls (Fig. 4). Putative methanol oxidation (*mxfA*) was most abundant in D1, D4, SW and WC, was present but in low abundance in D2 and D6, and was not identified in D3 (Fig. 4). While an in-depth interrogation of heterotrophic pathways is beyond the scope of this analysis, genes for fermentation were queried and found in all fluids, but at noticeably lower abundance in WC, SW and D1 than D2-6.

The capability of autotrophic carbon fixation is arguably an important one in carbon limited deep subsurface environments and is widespread at DeMMO. We detected genes for five of the six empirically demonstrated carbon fixation pathways (Hugler and Sievert, 2010) across samples. A list of genes queried, and which carbon fixation mode they are associated with can be found in Supplementary Data File 4. As illustrated in Fig. 5, genes indicative of the 3-hydroxypropionate bicycle were identified in MAGs from all 8 fluids. MAGs in D6 contained a relatively high abundance of *abfD*, a key gene in the 3-Hydroxypropionate/4-Hydroxybutyrate CO_2_ fixation cycle, but it was less common in other DeMMO fluids, and absent from WC and SW (Fig. 5). RuBisCO, which catalyzes the carboxylation and oxygenation of ribulose 1,5-bisphosphate (Tabita et al., 2008) was found at quite low abundances in all fluids, as were *aclA* and *aclB*, key genes in the reverse TCA cycle (Fig. 5). Importantly, genes for the Wood Ljungdahl pathway (*cooS, cdhE, cdhD*) were the most abundant in DeMMO fluids when compared to the other four CO_2_ fixation cycles queried, but were not detected in the controls, SW and WC (Fig. 5).

**Figure 5.**
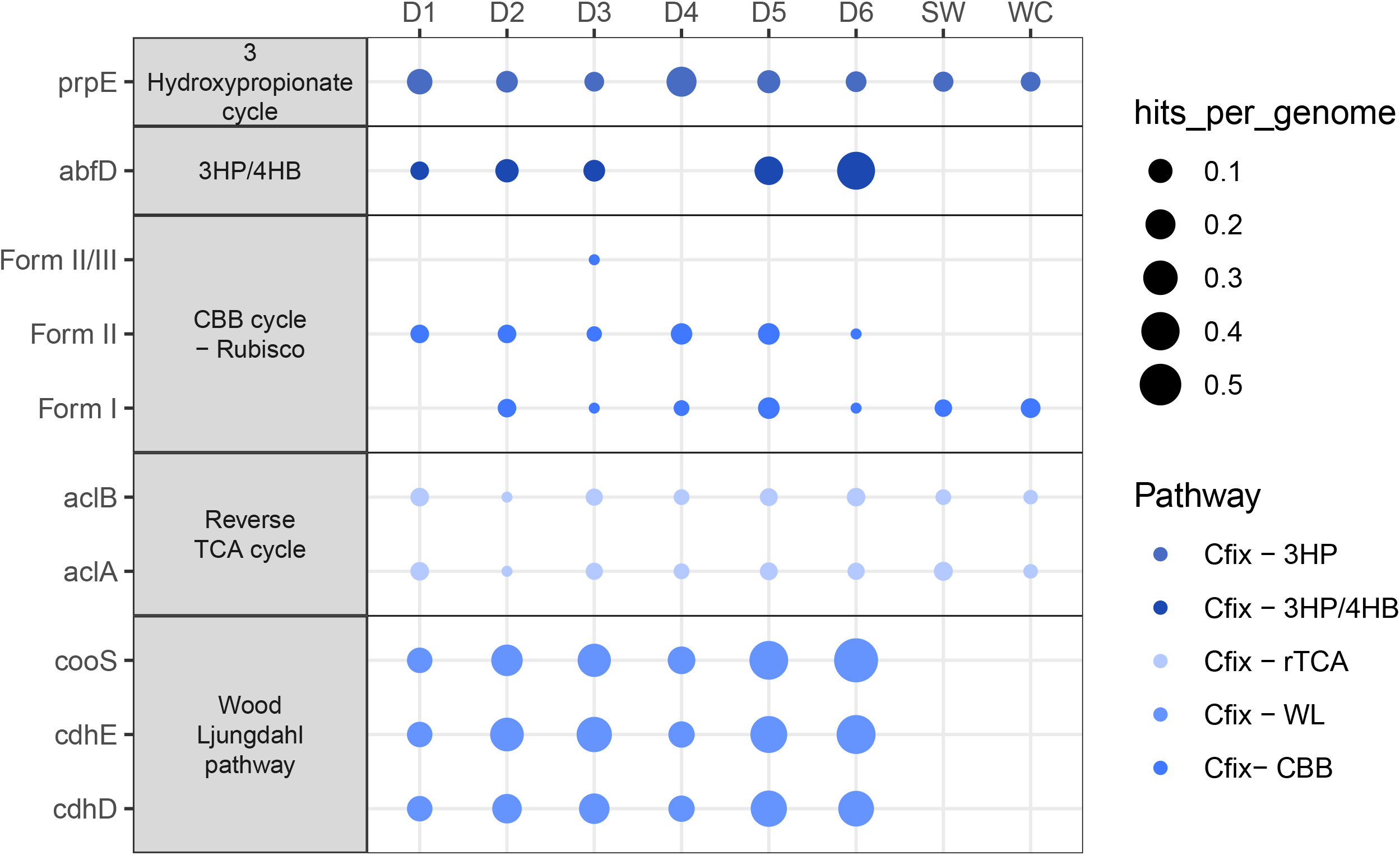
Modes of carbon fixation in MAGs, across all eight fluid samples. Five of the six documented modes of carbon fixation are included in this figure, for which we have deterministic, essential canonical marker genes. Gene abbreviation are listed to the right of the plot, along with full pathway name. Gene abundance was normalized by number of reconstructed MAGs for each site individually.

## DISCUSSION

### Controls and possible contamination

The control samples, WC and SW, were included in our analysis to assess their overall taxonomic and metabolic similarities and differences relative to DeMMO subsurface sites. We found stark differences in high-level taxonomic groups (e.g.-absence of Archaea from control samples, presence of Cyanobacteria in both control samples but not DeMMO fluids-Fig. 2). Differences in putative metabolisms between DeMMO and control fluids (e.g.-absence of hydrogenases and metal reducing genes and relatively high abundance of oxygen respiring genes in control samples, Fig. 4) were also apparent. In the service water, which is used as lubricant during boring activities and was used to drill D4-D6 in 2016, we detected a high abundance of Proteobacteria, compared to the DeMMO fluids. This water originates as municipal water for the city of Lead, South Dakota and is stored underground in tanks until it is used for lubricant or other uses in the underground mining facilities. While these are not quantitative measurements of contamination or lack thereof, these differences combined with analyses from other DeMMO datasets show that DeMMO fluids are distinct from laboratory controls, surface waters, and service water (Osburn et al., 2019; Casar et al., 2020; Rowe et al., 2020). Indeed, previous 16S rRNA gene sequencing performed over a 4-year timeseries within the same DeMMO boreholes demonstrated that the service water has a significantly different microbial community compared to the DeMMO fluids, and that there is no evidence for cross contamination (Osburn et al., 2020). Because of qualitative differences observed between sites in this metagenomic dataset and those shown between controls and DeMMO fluids using 16S rRNA analyses, we are confident that the trends we describe are not driven by contamination acquired either within the mine during sampling or during laboratory processing. That said, fluid paths within the subsurface environment have the potential to exchange microbes. For instance, WC and D1 are likely hydrologically connected as are and D4 and D5 (Osburn et al., 2019) and thus a natural connection in the microbiology is to be expected. We focus our subsequent discussion specifically on the bacterial groups from the DeMMO subsurface fluids.

### Overall metabolic capabilities in DeMMO subsurface fluids

One key question regarding the microbial ecology of the deep terrestrial subsurface is that of geochemical cycling of labile elements (e.g.-carbon, iron, nitrogen, sulfur), and how subsurface microbes interact with those elements in their bioavailable forms (considering both assimilatory and dissimilatory processes). With increasing access to paired geochemical and biological (both DNA and RNA) information from subsurface environments around the globe, we can begin to understand the scope of subsurface microbial biogeochemical cycling. Although we cannot determine the microbial processes that are occurring with metagenomes, we can make predictions about what processes the DBS community is capable of at the DeMMO locations that vary geologically, geochemically, and spatially.

### Metabolic capabilities by taxonomic group

One significant benefit to binning MAGs from metagenomic data is that metabolic pathways can be linked to discrete genome bins with accompanying taxonomy. Here we discuss a suite of 27 potential metabolic reactions (Supplementary Data File 5) and tie them to specific cultivated and uncultivated phyla (Fig. 6, Supplementary Figs. 1-15). In the following Results and Discussion we will refer to MAGs with the notation ‘SAMPLE_#,’ indicating the fluid sample from which they were reconstructed and the numerical rank of that MAG’s relative completeness among all MAGs from that single sample.

**Figure 6.**
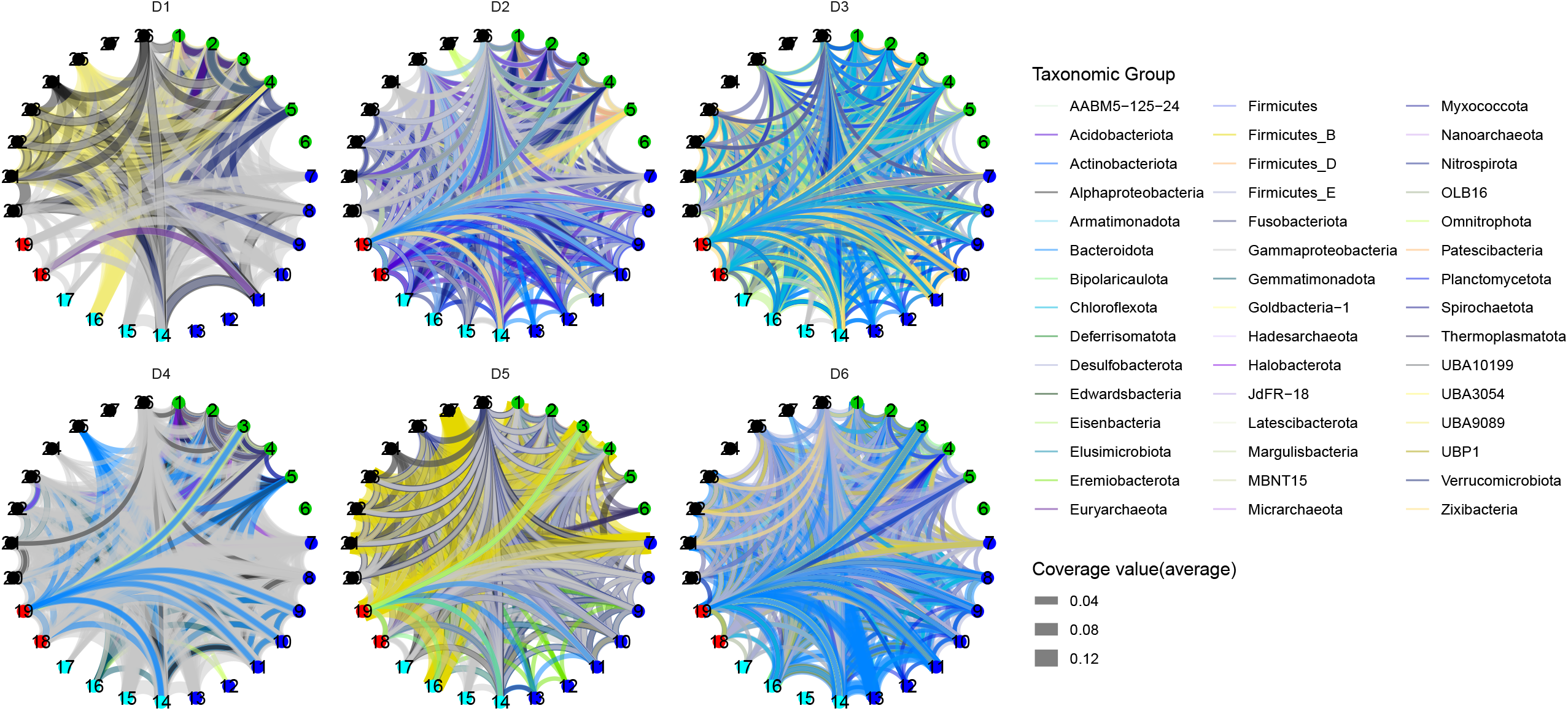
Metabolic chord diagrams for 27 energy yielding metabolic reactions of interest. Diagrams are separated by sample and represent the metabolic potential for all MAGs reconstructed from each respective fluid.

### Predicted metabolic capabilities of common cultivated phyla

Members of cultivated phyla likely play key roles in subsurface biogeochemical cycling at DeMMO. Members of the Proteobacteria have broad metabolic capabilities, mediating 25 of the 27 reactions that were queried, with the exception of methanogenesis (# 6) and sulfite reduction (# 27) (Supplementary Figs. 1-3). Nitrospira MAGs also have broad metabolic potential including for heterotrophy, nitrogen and sulfur metabolisms, but also metal reduction and hydrogen oxidation (Fig. 6, Supplementary Fig. 4). These observations are consistent with what is known from cultivated Nitrospira (Daims and Wagner, 2018).

Members of the Firmicutes have been found in global subsurface environments and often increase in abundance with depth and play key roles in S-cycling (Baker et al., 2003; Cowen et al., 2003; Chivian et al., 2008; Aüllo et al., 2013; Tiago and Veríssimo, 2013; Magnabosco et al., 2015; Jungbluth et al., 2016). Firmicutes (including the newly delineated Firmicutes groups B, D and E) are quite common in D6 (15 MAGs), and four MAGs each from Firmicutes group E and B were reconstructed from D3 and D5, respectively (Figure 2). Firmicutes in D6 have a wide array of potential metabolisms, including methanotrophy (#5), carbon fixation (#4), hydrogen generation (#19) and selenate reduction (#16) (Supplementary Figs. 5-7). Although there are only four Firmicutes group B MAGs identified in D5, they appear capable of diverse metabolisms involving all elements queried (carbon, hydrogen, nitrogen, oxygen, sulfur, metals) except for those dissimilatory metabolisms involving nitrogen species (Supplementary Fig. 6).

MAGs from the recently cultivated phylum Elusimicrobia (Geissinger et al., 2009) were identified in D3-D6 fluids. The MAG in D3 (DeMMO3_73) may be capable of fermentation and/or coupling carbon oxidation to selenate or arsenate reduction. Putative Elusimicrobia in D5 (DeMMO5_39 and _42) may be active in carbon and nitrogen cycling in those fluids, possibly capable of carbon fixation, simple carbon compound oxidation with selenate reduction, and possibly nitrite ammonification (#11). The single Elusimicrobia genome reconstructed from D6 fluids appears to be quite metabolically diverse although incomplete (50.0%/0.6% completeness/contamination) containing genes for reactions involving all elements of interest, save those involving hydrogen generation (#19) or oxidation (#18) (Supplementary Fig. 8).

### Predicted metabolic capabilities of uncultivated bacterial phyla

This dataset contains a wealth of MAGs from uncultivated bacterial groups. The ability to assign metabolic processes to members the candidate phyla is particularly powerful as it is one of the few sources of information on these abundant and under characterized organisms. This information can further assist in metagenome informed, targeted culturing of novel groups, adding to its value. For example, OLB16 is a recently named bacterial phylum, for which there are only two publicly available genomes (https://gtdb.ecogenomic.org/searches?s=al&q=OLB16, accessed October 10, 2020). One of these genomes was recovered from D6 fluids in a 2017 study (Momper et al., 2017). In this previous study we predicted that OLB16 (SURF_12) may be capable of nitrate reduction, sulfite oxidation, methane oxidation, and carbon fixation via the Wood Ljungdahl pathway (Momper et al., 2017). Here, we reconstructed two more OLB16 MAGs in D3 and D6 fluids (DeMMO3_89, DeMMO6_4). These new MAGs appear capable of sulfur oxidation and/or reduction (#21-23), heterotrophy (#1), fermentation, and metal reduction (#17). The more complete MAG in D6 also appears to transform hydrogen (#18, #19) (Supplementary Fig. 19), indicating a wide repertoire of potential metabolisms for this newly identified candidate phylum.

Additional newly identified phyla (Parks et al., 2017) were found in DeMMO fluids: UBA 9089, UBA3054, and UBA10199. The uncultivated phylum UBA9089 was identified in D1 and D3-D5 (Figure 2). Among those MAGs, sulfur and carbon cycling metabolisms are ubiquitous, with MAGs in D1, D4 and D5 potentially capable of selenate reduction (#16) and perhaps even capable of carbon fixation (#4) (Supplementary Fig. 10). UBA3054 (Parks et al., 2017), was identified in D6 fluids (DeMMO6_5). Currently there are only three publicly available genomes from this newly identified phylum, of which DeMMO6_5 is by far the most complete (other available MAGs are ∼54-58% compared to ∼99% complete). This MAG contains genes capable of heterotrophy (#1), fermentation (#3), arsenate reduction (#14), elemental sulfur oxidation (#21) and hydrogen generation (#19) (Supplementary Fig. 12). UBA10199 was found almost exclusively in D2 fluids (DeMMO2_63 and _70) with one MAG recovered from D3. These MAGs appear capable of heterotrophy (#1-3) and elemental sulfur (#21) and thiosulfate oxidation (#25) (Supplementary Fig. 12). The phylum UBP1, first reported in 2017 (Parks et al., 2017), was found in D5-6 (DeMMO5_30, DeMMO6_31). Metabolic reconstruction indicates that members of this phylum have diverse metabolic capabilities, potentially mediating sulfur reduction and/or oxidation (reactions #21-23), nitric oxide reduction (#10), and ammonification (#11) (Supplementary Fig. 13).

A single MAG from the candidate phylum Latescibacteria, formerly known as WS3, was found in D2 fluids (DeMMO2_58). Despite its relative incompleteness (∼48% complete), this MAG contains diverse metabolic potential with genes involved in 16 of the 27 reactions queried, most notably for H_2_ generation and oxidation and metal reduction (reactions #17-19) (Supplementary Fig. 14). Two MAGs from the recently designated candidate phylum Margulisbacteria (formerly candidate division ZB3) were found in D3 (DeMMO3_43 and _47). These appear to be capable of elemental sulfur and carbon oxidation coupled to arsenate reduction (#14) and potentially fermentation (Supplementary Fig. 15). Establishment of fermentation and hydrogen dependent metabolisms in this candidate phylum have been well documented (Carnevali et al., 2019; Utami et al., 2019), potential sulfur oxidation and arsenate reduction have not been reported, and may point to a differing ecological niche, due to the differing metabolic role for these new MAGs.

### Carbon fixation in DeMMO subsurface fluids

Although photosynthetically derived organic carbon does infiltrate into Earth’s subsurface, it is often recalcitrant and at very low concentrations (Pedersen, 2000). Dissolved organic carbon (DOC) concentrations at DeMMO sites are very low (generally <1 mg/L) (Osburn et al., 2019). As this DOC is limited and potentially of low quality, autotrophic carbon fixation may be a key metabolic adaptation, especially in the deeper, older DeMMO fluids. Quite telling is the fact that genes involved in the Wood Ljungdahl pathway were by far most abundant across all six DeMMO fluids yet not detected in the controls (Figure 5). This trend is supported by the geochemical conditions in these fluids (Table 1). At the time of sampling, the oxidation reduction potential in DeMMO fluids ranged from -61 to -198 mV, indicating little to no oxygen present. The Wood Ljungdahl pathway requires anoxic conditions, as some of its enzymes, especially the crucial acetyl-CoA synthase, are highly oxygen sensitive (Berg, 2011). In contrast, the SW and WC fluids are at near oxygen saturation (∼300 mV, data not shown), making it highly unlikely this form of carbon fixation could be performed. The Wood Ljungdahl pathway has previously been hypothesized to be a crucial source of primary production in D6 (Momper et al., 2017) and other deep terrestrial subsurface environments (Magnabosco et al, 2015), a conclusion supported by this study. Similarly, it follows that RuBisCo, the key enzyme which catalyzes the carboxylation and oxygenation of ribulose 1,5-bisphosphate (Tabita et al., 2008) was not common in the dark and reducing fluids of DeMMO (Figure 6). Other trends in carbon fixation gene presence are not as straightforward. For example, the 3-hydroxypropionate bi-cycle is commonly used by members of the Chloroflexi (Hügler and Sievert, 2010), yet the fluids with the greatest number of Chloroflexi MAGs (D6) do not have the greatest proportion of identified canonical *prpE* genes. Along the same vein, the 3-Hydroxypropionate/4-Hydroxybutyrate mode of carbon fixation is typically associated with the archaea (Berg et al., 2007; Loder et al., 2016), yet in this study we find the highest proportion of *abfD* genes in MAGs from D6 (Figure 6), fluids from which we did not construct archaeal MAGs (Figure 2).

### Tying metabolisms to in situ geochemistry

Combining geochemistry with microbiology is challenging in any environmental setting, but is especially demanding in deep, difficult to access, subsurface environments. Monitoring at the DeMMO observatory has enabled a rare opportunity to combine 16S rRNA tag sequencing, whole metagenomic DNA sequencing and assemblies, and geochemical analyses over a multi-year period. Other terrestrial subsurface studies have combined geochemical analyses with extensive metagenomic datasets to gain insight into elemental cycling, microbial community connectedness and microbial metabolic ‘handoffs’ (Anantharaman et al., 2016; Lau et al., 2014; Magnabosco et al., 2016). These studies found that deep (> 1 km) subsurface communities tend to rely heavily on the Wood Ljungdahl pathway for potential carbon fixation and primary production (Magnabosco et al., 2016) and that some subsurface habitats seem to preserve ancestral gene signatures in biogeochemically relevant functional genes (Lau et al., 2014).

In general terms, the distribution of metabolically functional genes identified in this MAG dataset agree well with our understanding of *in situ* geochemical conditions at DeMMO. As detailed in the results, we find broad metabolic potential for biogeochemical cycling across the DeMMO sites, but variation both between sites. For example, sulfate and nitrate are available TEAs in all fluids, and sulfur and nitrogen cycling genes are abundant across all DeMMO samples (Table 1 and Fig. 4), particularly those sites where there are abundant forms of N and S (D4, D5). Along these same lines, hydrogenases were identified in abundance in D2-6, but were far less abundant in D1, the DeMMO borehole with the lowest average dissolved hydrogen concentration, and were nearly or completely absent from MAGs binned from the oxic control samples (Table 1 and Fig. 4).

However, deeper examination of the MAGs in this DeMMO dataset revealed a number of instances where the most abundant functional genes in a given site were not consistent with the most abundant geochemical species at the time of DNA collection 2018, or in previous sampling campaigns over the last 4 years. For example, D4 has historically extremely low dissolved oxygen levels (∼0.03 mg/L) and during sampling in 2018 the measured ORP was -200 mV, however, we identified a wide variety and relatively large abundance of oxygen respiring genes in D4 MAGs (Table 1 and Fig. 4). It is therefore not clear where these organisms are acquiring the oxygen necessary for aerobic respiration. In terms of carbon metabolisms, genes involved in C1 metabolism and fermentation have varying patterns across all DeMMO fluids, but are quite abundant in all samples (Fig. 4). But dissolved organic carbon levels are quite low in all fluids (<0.5 and <3.0e^-6^ mg/L, respectively).

In general, the average gene count for MAGs recovered from DeMMO boreholes was not small, at ∼1,900-2,400 genes per MAG, with typical larger genomes containing ∼4,000-6,000 genes (Supplementary Data File 6). This leads us to question why deep subsurface microbes would retain genes and entire functional pathways when the geochemical substrate for that metabolism is not present or present at such low concentrations that the reaction is not energetically favorable. We hypothesize that microbes in this deep subsurface environment gain a competitive edge by maintaining a wide variety of functional pathways. Indeed, in this study we identified diverse putative metabolisms in MAGs from little-studied groups such as the Elusimcrobia and candidate phyla such as OLB16. Our findings indicate that the capability to perform many dissimilatory energy metabolisms is the norm rather than the exception in non-CPR MAGs in this deep subsurface environment. Other groups have reported similar findings in shallow and deep subsurface environments. Large, robust genomes were found in anoxic/near-anoxic and TEA-limited shallow aquifer environments (Anantharaman et al., 2016). When electron acceptors such as oxygen and nitrate were injected into these aquifers, there was an almost instantaneous draw down of those substrates, and they were again near detection limit within hours (Anantharaman et al., 2016). Similarly, the deep subsurface isolate *Desulforudis audaxviator* contains the genomic potential for an astonishing array of catabolic and anabolic metabolisms including sulfate reduction, carbon and nitrogen fixation, heterotrophy, and others (Chivian et al., 2008). Our findings support the growing body of evidence that in many subsurface biomes there is a dichotomy of small, ultra-streamlined genomes and larger, bulky genomes with diverse metabolic capabilities (Anantharaman et al., 2016; Jungbluth et al., 2016; Jungbluth et al., 2017; Lau *et al*., 2016).

With time series data (Osburn et al., 2019), long term *in situ* experiments (Casar et al., 2020) and metagenomic surveys (Momper et al., 2017, this study) we are beginning to piece together how planktonic and attached microbial communities vary temporally and interact with and influence *in situ* geology and geochemistry in a deep terrestrial subsurface environment, DeMMO. In addition to long-term, incremental understanding about terrestrial subsurface processes, metagenomic studies such as these can significantly augment our understanding of novel microbial groups and their role in these environments. This understanding enables deeper questions, such as what percentage of elemental cycling can be attributed to biotic vs abiotic processes, and on what timescales these processes occur for the whole of the DSB and for environments that may be of particular interest. While no two subsurface sites are the same, there are commonalities including an enrichment of Firmicutes, uncultured candidate phyla, and abundant members of the CPR. Metabolically these groups have the potential for widespread carbon utilization and fixation, sulfur and metal-based metabolisms, and potential roles in rapid drawdown of injected TEAs such as nitrate and oxygen.

## CONCLUSIONS

A pillar of advancing environmental microbiology rests on culture independent methods to assessment of metabolic potential of microbes. Here we present nearly 600 high quality MAGs deriving from an established transect through the subsurface, from the surface to 1.5 km deep into the crust. This dataset includes a diversity of taxonomic groups including those from well-cultivated but metabolically important lineages like the Proteobacteria and Nitrospira, as well as from virtually unknown taxa known only from a small number of assembled genomes. In these genomes we find the genetic potential to mediate a range of environmentally important metabolisms including nitrogen, sulfur, and metal cycling as well as C1-metabolisms, fermentation, and carbon fixation. The spatial distribution of this metabolic potential often agrees well with available chemical species, but there are intriguing instances of disagreement which suggest that maintaining genomic plasticity is a key adaptive strategy of many intraterrestrial microbes. This work fits into a growing body of work at DeMMO which facilities integrated understanding of the deep subsurface biosphere in space, through time, and in chemical and environmental context. Further, these genomes add to and expand the growing body of subsurface genomes, informing the capabilities of novel taxa and expanding our ability to understand this biome on the global scale.

## DATA DEPOSIT

Sequence data for metagenomic assemblies and their respective MAGs and corresponding metadata can be accessed using the BioProject identifier PRJNA563685 and BioSample accessions SAMN18064095, SAMN18064236, SAMN18064310, SAMN18064413, SAMN18064496, SAMN18064575, SAMN18004502, and SAMN18005272 corresponding to sites DeMMO1-6, Whitewood Creek, and Service Water communities, respectively. All code and corresponding data used in this study are available at https://github.com/CaitlinCasar/Momper2021_DeMMO_FractureFluidMetagenomes.

## ACKNOWLEDGEMENTS

This work was supported by NASA Exobiology (NNH14ZDA001N) and grants to MRO from the David and Lucille Packard Foundation and the Canadian Institute for the Advancement of Research Earth 4D. We would like to recognize Michael D. Lee for helpful conversations on phylogenomic analyses, phylogenetic tree building and assistance in using GToTree. We also want to thank the developers of METABOLIC and Karthik Anantharaman for their help in running and interpreting the data generated using that package. We want to thank especially staff and personnel at SURF for access to the deep subsurface and repeated access to samples used in this study and Dr. Brittany Kruger of the Desert Research Institute for coordination of field sampling expeditions.

## SUPPLEMENTAL INFORMATION

Supplementary information is available at ISMEJ’s website

